# Sex-Specific Computational Models of Kidney Function in Patients with Diabetes

**DOI:** 10.1101/2021.07.17.452799

**Authors:** Sangita Swapnasrita, Aurélie Carlier, Anita T. Layton

## Abstract

The kidney plays an essential role in homeostasis, accomplished through the regulation of pH, electrolytes and fluids, by the building blocks of the kidney, the nephrons. One of the important markers of the proper functioning of a kidney is the glomerular filtration rate. Diabetes is characterised by an enlargement of the glomerular and tubular size of the kidney, affecting the afferent and efferent arteriole resistance and hemodynamics, ultimately leading to chronic kidney disease. We postulate that the diabetes-induced changes in kidney may exhibit significant sex differences as the distribution of renal transporters along the nephron may be markedly different between women and men, as recently shown in rodents. The goals of this study are to (i) analyze how kidney function is altered in male and female patients with diabetes, and (ii) assess the renal effects, in women and men, of an anti-hyperglycemic therapy that inhibits the sodium-glucose cotransporter 2 (SGLT2) in the proximal convoluted tubules. To accomplish these goals, we have developed computational models of kidney function, separate for male and female patients with diabetes. The simulation results indicate that diabetes enhances Na^+^ transport, especially along the proximal tubules and thick ascending limbs, to similar extents in male and female patients, which can be explained by the diabetes-induced increase in glomerular filtration rate. Additionally, we conducted simulations to study the effects of diabetes and SGLT2 inhibition on solute and water transport along the nephrons. Model simulations also suggest that SGLT2 inhibition, which constricts the afferent arteriole to attenuate glomerular hyperfiltration, can then limit Na^+^-glucose transport, consequently raising luminal [Cl^-^] at the macula densa and finally restoring the tubuloglomerular feedback signal. By inducing osmotic diuresis in the proximal tubules, SGLT2 inhibition reduces paracellular transport, eventually leading to diuresis and natriuresis. Those effects on urinary excretion are blunted in women, in part due to their higher distal transport capacity.

## 1 Introduction

In recent years, the role of sex and gender has emerged as a priority area in biological and medical research. In particular, there has been increasing evidence that sex has a significant impact on the pathogenesis of metabolic disorders, such as type 2 diabetes (T2D). Diabetes is currently estimated to be prevalent in 9.3% (463 million people) and expected to reach 10.2% (578 million) by 2030 and 10.9% (700 million) by 2045 [1]. In developed countries, T2D is associated with chronic kidney disease [2] and increases the risk of cardiovascular disease [3]. Interestingly, sex-specific differences have been reported in the disease prevalence and incidence of diabetes and diabetic kidney disease. Overall, men are predisposed at a higher rate to T2D and a similar prevalence of type 1 diabetes (T1D) compared with premenopausal women. Postmenopausal women, however, have an increased risk of developing glomerular hyperfiltration, diabetic kidney disease, and end-stage kidney disease, compared to age-matched men [4, 5].

To understand the origin of the differences in the prevalence in diabetes-induced renal complications between men and women, one must first understand kidney function and its sex differences. The kidney’s nephrons are the centers for filtration of electrolytes and water in the blood and the maintenance of pH. Each human kidney contains about a million nephrons, which are linked to clusters of glomerular capillaries that receive blood from individual afferent arterioles branching off into intra-renal arteries. A portion of that blood passes through the glomeruli and enters the nephron. Along the different segments of the nephrons (namely, the proximal tubules, the loops of Henle, the distal convoluted tubules, the connecting tubules, the collecting ducts), the content of the tubular filtrate undergoes major changes, via epithelial transporter-mediated reabsorption or secretion of solutes, and reabsorption of water, to become urine. In both women and men, urine output matches fluid and solute intake as well as waste production. However, major sex differences may be observed in the epithelial transport processes of the kidney.

Sex hormones regulate nearly every tissue and organ, including the kidney [6–10]. Veiras et al. [11] reported sexually dimorphic patterns in the distribution of renal ransporters (electrolyte, channels, claudins) across the different nephron segments in male and female rodents. Their findings demonstrated that, female rat nephrons exhibit lower reabsorption of Na^+^, bicarbonates, and water along the proximal tubules, resulting in a downstream shift of the Na^+^ and water reabsorption in female kidneys. Along the distal nephron segments, the female kidney exhibits a higher abundance of key Na^+^ transporters, relative to male, resulting in similar urine excretion between the sexes. In addition, experimental evidence has shown that uremic toxin transporters also display sex differences [12–14], for example the organic anionic transporter (OAT)3 is expressed less in male than female mice, and the opposite is valid for OAT1 [15]. Our modelling studies in the male and female rat kidney have previously identified the functional implications of these molecular differences in renal handling of water, electrolytes, and glucose [16–18] and in renal nitric oxide bioavailability and medullary oxygenation [19–23]. For example, due to their lower Na^+^ activity in the proximal tubule, female rats are able to more rapidly excrete a saline load [16].

Some of the sex differences in rodent kidney function may translate to humans. Despite obvious inter-species differences, women and female rats face similar challenges of circulating volume adaptation during pregnancy and lactation. In a recent study, we developed sex-specific models of epithelial transport along the nephrons in the kidney of a man and a woman [24]. Model simulations indicate that sexual dimorphism in renal transporter patterns similar to that reported in rodents may better prepare women for the heightened demands on the kidney during pregnancy and lactation [24]. An important open question remains: How do these findings translate to kidney function in diabetes? Indeed, differences have been reported in the prevalence and severity of diabetic kidney disease between women and men [25]. Even though the pathogenesis of diabetic kidney disease remains incompletely understood, pathophysiological changes that diabetes induces in the kidneys have been characterized. At the very onset of diabetes, the kidney enlarges and glomerular filtration rate (GFR) becomes supranormal [26]. Moreover, diabetes induces an alteration in transporter expression: the activity of SGLT2, GLUT2, NKCC and Na/K-ATPase are upregulated whereas the SGLT1 activity is downregulated, and this in a nephron segment specific manner [27]. These structural and hemodynamics changes affect kidney function and may eventually lead to chronic kidney disease. How might kidney function decline differ in male and female patients with diabetes, and why?

A related issue concerns the anti-hyperglycaemic drugs that target the kidney. As mentioned above, the kidney plays a major role in homeostasis, including the regulation of blood glucose levels [28]. Following glomerular filtration, almost all glucose is reabsorbed from the lumen of the kidney within the proximal tubule, via two major Na^+^-glucose cotransporters (SGLT1 and SGLT2), such that there is no loss of glucose. When insulin production is reduced, blood glucose levels increase and the task of handling this excess glucose load falls on the kidney. In patients with diabetes, filtered blood glucose levels exceed the transport capacity of SGLT1 and SGLT2, leading to the excretion of glucose in urine, i.e., glycosuria.. Historically, glycosuria has been associated with diabetes, but with the prescription of SGLT inhibitors, this pathophysiological condition is now used as a mechanism to lower blood glucose [29, 30]. Specifically, SGLT2 inhibitors block the glucose reabsorption along the early proximal tubule and induce glycosuria to reduce blood glucose levels. Since SGLT2 mediates the co-transport of glucose and Na^+^, inhibition of SGLT2 induces natriuresis and diuresis upon reduction of Na^+^ and fluid reabsorption in the proximal tubule. Thus, besides their intended anti-hyperglycaemic effect, SGLT2 inhibitors have been shown to lower blood pressure and provide protection from heart failure [31, 32]. Given how the promises of SGLT2 inhibitors, it seems imperative to thorough understand their mechanisms of action, some of which have remained unclear, as well as any sex-dependent kidney response to these drugs.

We have previously conducted model simulations to investigate kidney function in diabetes and the renal effects of SGLT2 inhibition on the kidney of a male rat or human [27, 33–35]. As such, even though some of the model predictions may generalize to women, the translational value of any study that involves only half of the population remains limited. Thus, the goal of the present study is to develop sex-specific models that allow us to analyze and compare kidney function in male and female patients with different stages of diabetes, and the renal effects of SGLT2 inhibition.

## 2 Materials and Methods

We previously implemented an epithelial cell-based model of transporter-mediated solute transport along the nephrons of a human kidney [24, 36]; that model was recently extended to simulate kidney function in a male patient with diabetes [33]. In this study, we extend that model to simulate kidney function in a female patient with diabetes. The model represents the following six classes of nephrons: a superficial nephron, which turns at the outer-inner medullary boundary, and five juxtamedullary nephrons, of different lengths, each reaching into differing levels of the inner medulla. While the superficial nephrons account for 85% of the nephron population, and extend from the Bowman’s capsule to the papillary tip, the remaining 15% of the nephrons are juxtamedullary that possess loops of Henle that reach to different depths in the inner medulla; most of the long loops turn within the upper inner medulla. Population-based weighted average is taken wherever necessary. Each model nephron is represented as a tubule with a variety of transporters on the apical and basolateral membrane. The model assumes that ten connecting tubules coalesce successively to one cortical collecting tubule [37]. In the inner medulla, the collecting ducts again coalesce successively in the ratio of 10:1. A schematic diagram for the model is shown in Fig. 1.

**Figure 1.**
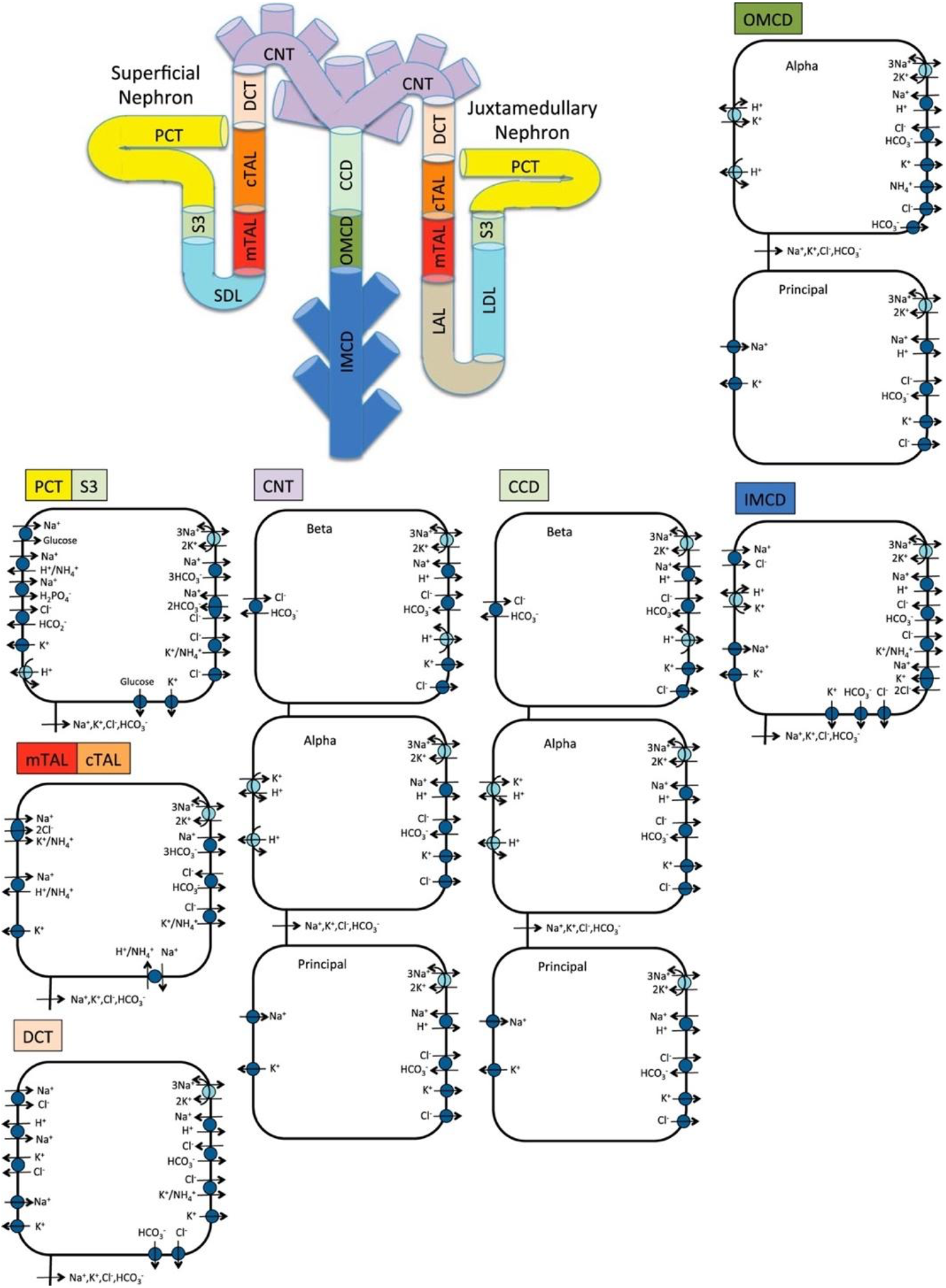
Schematic diagram of the nephron system (not to scale). The model includes 1 representative superficial nephron and 5 representative juxtamedullary nephrons, each scaled by the appropriate population ratio. Only the superficial nephron and one juxtamedullary nephron are shown. Along each nephron, the model accounts for the transport of water and 15 solutes (see text). The diagram displays only the main Na^+^, K^+^, and Cl^−^ transporters. mTAL, medullary thick ascending limb; cTAL, cortical thick ascending limb; DCT, distal convoluted tubule; PCT, proximal convoluted tubule; CNT, connecting duct; CCD, cortical collecting duct; SDL, short or outer-medullary descending limb; LDL/LAL, thin descending/ascending limb; OMCD, outer-medullary collecting duct; IMCD, inner-medullary collecting duct.

The model accounts for the following 15 solutes: Na^+^, K^+^, Cl^−^, HCO_3_^−^, H_2_CO_3_, CO_2_, NH_3_, NH_4_^+^, H_2_PO_4_^−^, H^+^, HCO_2_^−^, H_2_CO_2_, urea, and glucose. The model is formulated for steady state and consists of a large system of coupled ordinary differential equations and algebraic equations [36]. Model solution describes luminal fluid flow, luminal fluid solute concentrations, and hydrostatic pressure. Excluding the descending limb segment, model solution also describes cytosolic solute concentrations, membrane potential, and transcellular and paracellular fluxes. In a non-diabetic kidney, we assume a single-nephron glomerular filtration rate (SNGFR) of 100 and 133 nl/min for the superficial and juxtamedullary nephrons, respectively, in both women and men. Assuming a total of 1 million nephrons, this yields a GFR of 105 mL/min. Model parameters that describe a non-diabetic human kidney can be found in [36].

### 2.1 Sodium-glucose cotransport in the proximal tubule

Under euglyceamic conditions, the SGLTs facilitate the reabsorption of most of the filtered load of glucose, working in tandeom with the glucose transport facilitators (GLUT) on the basolateral side of the proximal tubule cells. The proximal convoluted tubule expresses the high-capacity, low-affinity transporter SGLT2 and GLUT2.; the S3 segment expresses the lower capacity, higher affinity transporter SGLT1 and GLUT1. The modeling of glucose and Na^+^ fluxes across SGLT2 and SGLT1 cotransporters, and glucose fluxes across GLUT1 and GLUT2 have been described in our previous studies [27, 35, 38, 39].

### 2.2 Simulating sex-specific kidney function

We have previously developed sex-specific models of kidney function for the rat [16, 17] and for humans [24]. To produce those models, we formulated epithelial cell-based models of solute transport along male and female rat nephrons. First, sex-specific epithelial transport models were formulated only for the proximal tubule, then for the thick ascending limb, for the distal convoluted tubule, for the connecting tubule, and then individually for the cortical and medullary segments of the collecting ducts. The male and female transport models account for sex differences via the expression levels of apical and basolateral transporters [11, 40]; which also vary among nephron segments. Nephron segment lengths and luminal diameters in the human kidney are taken to be the same in women and men due to the absence of sex-specific human data; additional model parameters can be found in [36]. Key differences in nephron transport parameters between non-diabetic women and men are summarized in Table S1 (Supplementary materials).

### 2.3 Simulating a diabetic kidney

Diabetes is associated with renal hypertrophy, hyperfiltration, and alterations in transporter expression [4, 5, 28, 41]. In this study, we simulate two diabetic conditions:

1. moderate diabetes: plasma glucose is elevated from its non-diabetic value of 5 mM to 8.6 mM; SNGFR is increased by 27% [26]; the tubular diameter and length of the proximal tubules are increased by 10%; and the diameter and length of the distal segments are increased by 18 and 7% [27, 42];
2. severe diabetes: plasma glucose is further increased to 20 mM; SNGFR is increased by 10% [26]; the tubular diameter and length of the proximal tubules are increased by 28%; the diameter and length of the distal segments are increased by 42 and 7%, respectively [27, 42].

Due to lack of sex-specific data, we assume the same morphological changes in both men and women and a similar enhancement in transcellular water permeability along the cortical and inner-medullary collecting duct segments by 55 and 40%, respectively, in moderate and severe diabetes [27]. The altered transporter activities are summarized in Table 1.

**Table 1.**
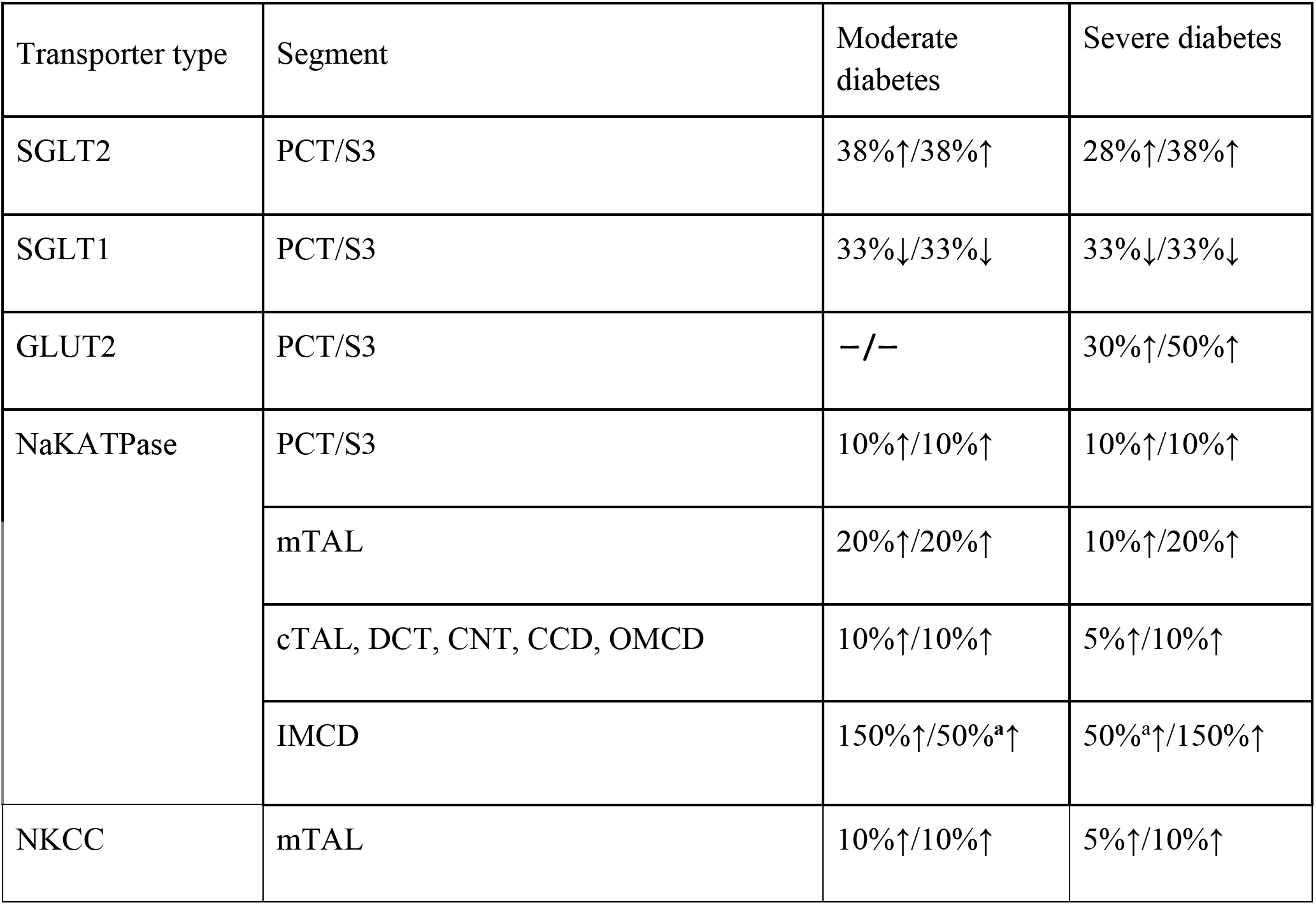
Up (↑)- or downregulation (↓) of transporter activity as [female value]/[male value] along different nephron segments in moderate and severe diabetes. PCT, proximal convoluted tubule; mTAL/cTAL, medullary/cortical thick ascending limb; DCT, distal convoluted tubule; CNT, connecting tubule; CCD/OMCD/IMCD, cortical/outer-medullary/inner-medullary collecting duct. ^**a**^50% along the first 2/3rd of the segment and 150% along the rest.

| Transporter type | Segment | Moderate diabetes | Severe diabetes |
| --- | --- | --- | --- |
| SGLT2 | PCT/S3 | 38%↑/38%↑ | 28%↑/38%↑ |
| SGLT1 | PCT/S3 | 33%↓/33%↓ | 33%↓/33%↓ |
| GLUT2 | PCT/S3 | –/– | 30%↑/50%↑ |
| NaKATPase | PCT/S3 | 10%↑/10%↑ | 10%↑/10%↑ |
|  | mTAL | 20%↑/20%↑ | 10%↑/20%↑ |
|  | cTAL, DCT, CNT, CCD, OMCD | 10%↑/10%↑ | 5%↑/10%↑ |
|  | IMCD | 150%↑/50%<sup>a</sup>↑ | 50%<sup>a</sup>↑/150%↑ |
| NKCC | mTAL | 10%↑/10%↑ | 5%↑/10%↑ |

### 2.4 Simulating SGLT2 inhibition

We assume that following acute SGLT2 inhibition, SNGFR decreases by 3% in all nephrons, in accordance to the minor 3% GFR reduction seen in non-diabetic subjects, being administered canagliflozin or dapagliflozin for 4 days [27]. Also, in non-diabetic subjects, SGLT2 inhibition yields a higher urinary excretion of glucose (45% of filtered load) [43]. In our previous study, we modelled 90% inhibition of SGLT2 in all nephrons, resulting in the excretion of 40% of the filtered glucose, to approximate this glucose excretion [43]. SGLT2 inhibition attenuates diabetic-reduced hyperfiltration [44]. Thus, when simulating the effects of SGLT2 blockade in a diabetic kidney, we lower GFR to its non-diabetic level of 151.2 L·day^−1^. Upon acute administration, SGLT2 inhibition is assumed not to affect plasma glucose concentration; thus, blood glucose level is kept at 8.6 and 20 mM in the moderate and severe diabetes cases, respectively.

## 3 Results

### 3.1 Kidney function under diabetic conditions

We investigate the change in solute and water transport along the nephrons due to diabetes, and if those changes differ between men and women. Key results are summarized in Figs. 2–4. In these simulations, we mimic the renal effects of moderate and severe diabetes as described in the Materials and Methods. In particular, diabetes induces glomerular hyperfiltration and tubular hypertrophy, to different extents depending on the severity of the disease. Elevated GFR is reflected on filtered solute loads, whereas tubular hypertrophy results in enhanced transport, as discussed below. We will first compare nephron transport in a non-diabetic and diabetic kidney between the two sexes. Then we will dive deeper into the sex differences.

**Figure 2.**
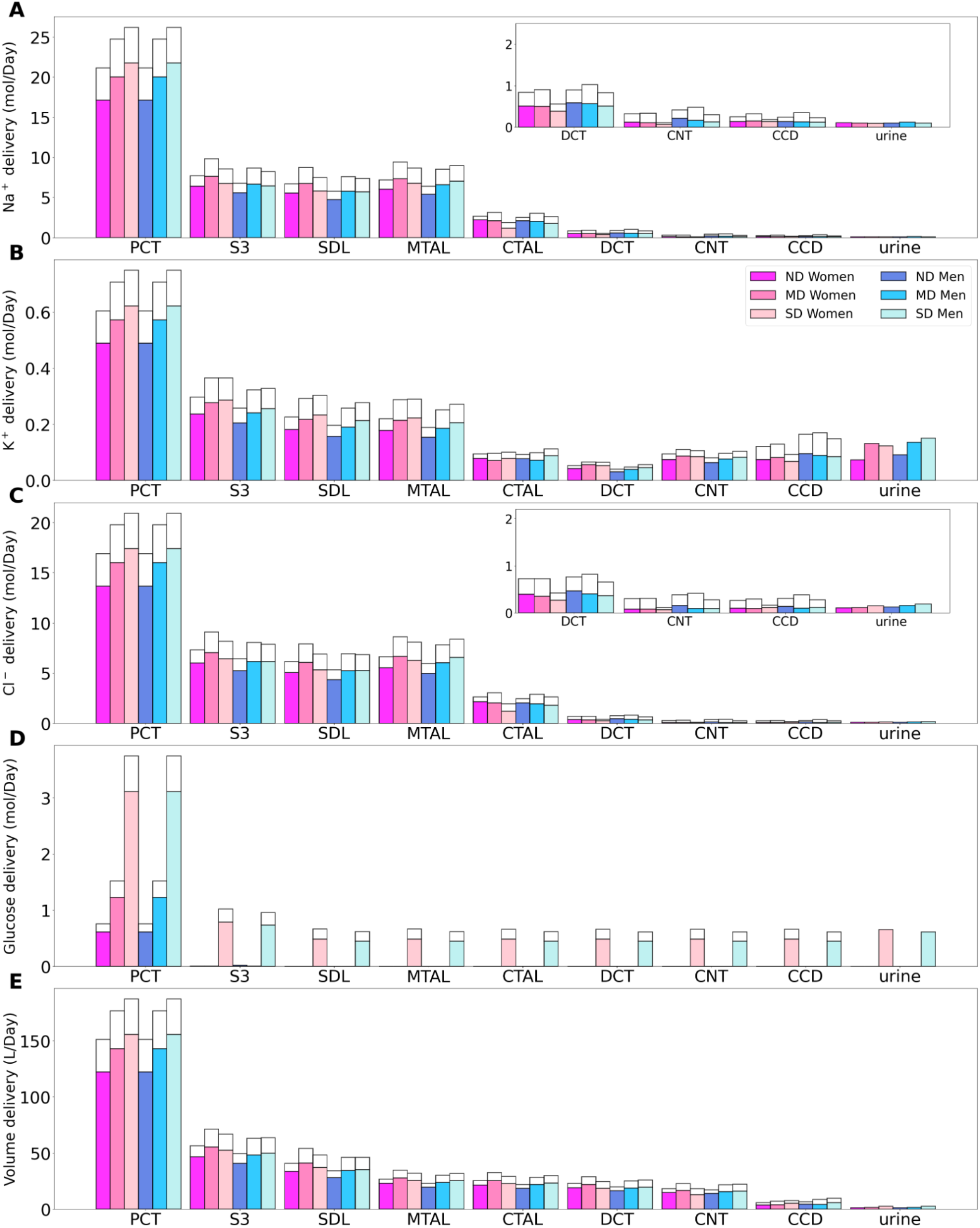
Delivery of Na^+^ (*A*), K^+^ (*B*), Cl^-^ (*C*), glucose (*D*) and fluid (*E*) to the beginning of individual nephron segments in women and men without diabetes (ND) and with moderate (MD) and severe diabetes (SD). Color bars, superficial nephron values; white bars, juxtamedullary values, computed as weighted totals of the five representative model juxtamedullary nephrons. The models assume that superficial nephrons account for 85% of the nephron population; thus, the superficial delivery values are substantially higher. In each case, only one bar is shown for “urine” since the superficial and juxtamedullary nephrons have merged at the cortical collecting duct entrance. PCT, proximal convoluted tubule; SDL, descending limb; MTAL/CTAL, medullary/cortical thick ascending limb; DCT, distal convoluted tubule; CNT, connecting tubule; CCD, cortical collecting duct.

**Figure 3.**
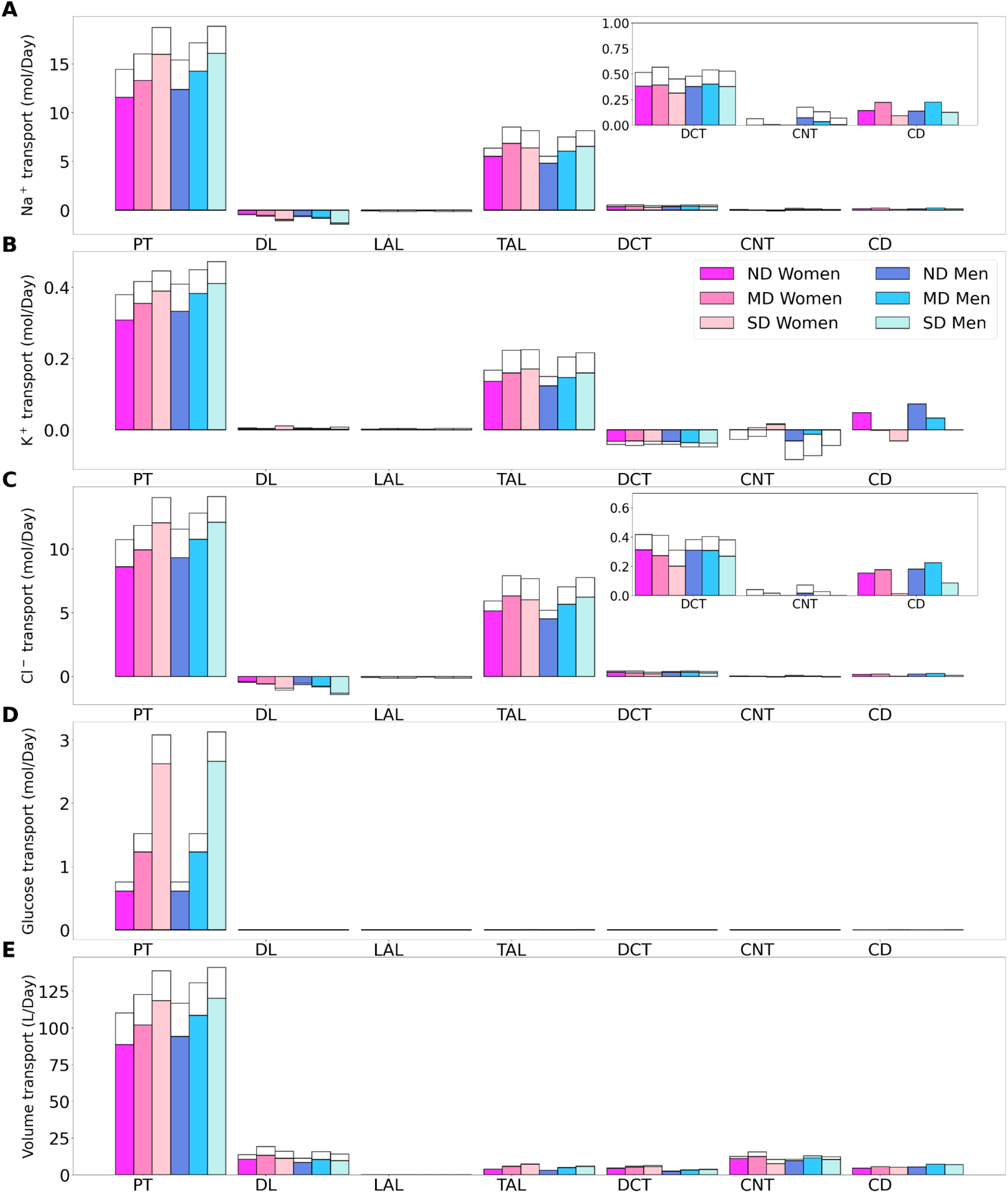
Transport of Na^+^ (*A*), K^+^ (*B*), Cl^-^ (*C*), glucose (*D*) and fluid (*E*) along individual nephron segments in women and men without diabetes (ND), with moderate diabetes (MD), and with severe diabetes (SD). Notations are analogous to Fig. 2. PT, proximal convoluted tubule and S3.

**Figure 4.**
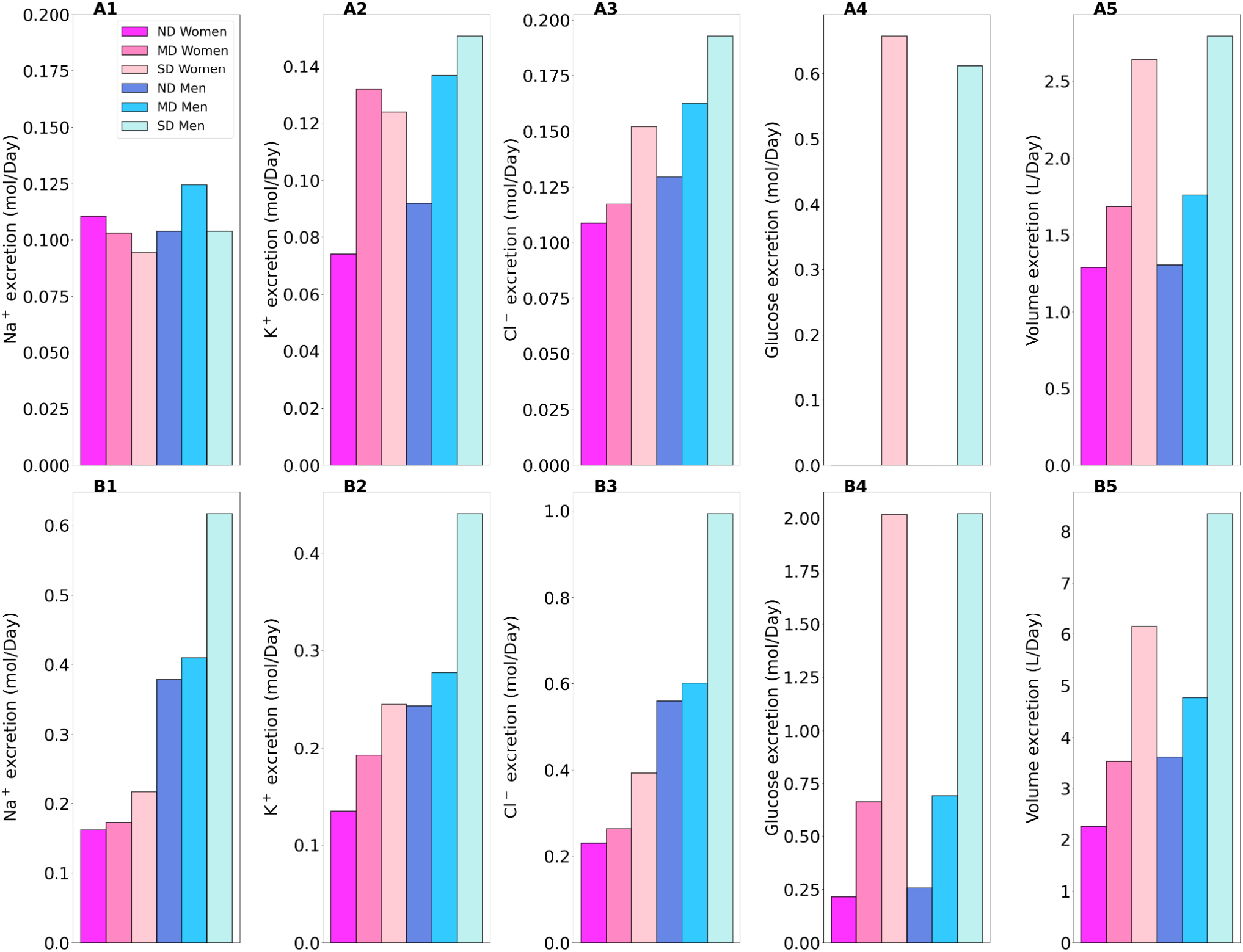
Total excretion of Na^+^ (*1*), K^+^ (*2*), Cl^-^ (*3*), glucose (*4*) and fluid (*5*) in men and women without diabetes (ND), with moderate diabetes (MD), and with severe diabetes (SD). Results are obtained under no inhibition (*A1-A5*) and with SGLT2 inhibition (*B1-B5*).

The proximal tubules in a healthy kidney reabsorb essentially all filtered glucose. In a healthy kidney, the proximal convoluted tubule in both women and men reabsorbs 97% of filtered glucose via the SGLT2, and the S3 segments reabsorb almost all of the remaining glucose via the SGLT1 (See Fig. 2D). In a diabetic kidney, the filtered glucose load increases substantially, but so does the glucose transport capacity. In the moderate diabetes simulations, plasma glucose concentration is assumed to increase from 5 to 8.6 mM, which together with the elevated GFR yields a filtered load of 1.52 mol·day^−1^glucose, which is twice the regular filtered load. However, as the proximal tubules hypertrophize under diabetic conditions, they gain10% in both length and diameter, which corresponds to ~20% increase in effective transport area. Thus, the enhanced proximal tubule glucose transport balances the higher filtered load, resulting in the absence of glycosuria (glucose excretion in urine) in patients with moderate diabetes [41].

In severe diabetes simulations, plasma glucose is further elevated to 20 mM, resulting in glucose filtered load of 3.75 mol·day^−1^ glucose. The length and diameter of the proximal tubules further increase to 28% above baseline, resulting in a 64% increase in effective transport area. Despite its enhanced transport capacity, the glucose load exceeds the transport capability of the proximal tubule. Net glucose reabsorption by the proximal convoluted tubule and S3 segment increases, but because of the increased filtered load, fractional reabsorption along the proximal convoluted tubule is predicted to decrease to 72.7% in women and 74.5% in men, whereas the S3 accounts for 0.1% of the filtered glucose in both sexes. Because downstream segments do not possess significant glucose transport capacities. The severe diabetic model predicts female and male glucose excretion of 0.65 and 0.61 mol·day^−1^, respectively.

Besides glucose, diabetes affects renal transport of electrolytes and water as well. The 10% increase in GFR in the moderate diabetic kidney, relative to a healthy kidney, implies a 10% increase in filtered Na^+^ and also ~10% increase in total Na^+^ transport, since only ~1% of the filtered Na^+^ is excreted. The model predicts that the increased transport activities occur primarily by the tubular segments where the Na^+^ transporter activities are significantly elevated in diabetes (Fig. 3A). In particular in the proximal convoluted tubules the hyperfiltration-induced changes in the torque augment the density of all transcellular transporters, and in the medullary thick ascending limbs diabetes enhances the density of NKCC2 [27, 45]. Then, enhanced Na^+^ transport can compensate for the elevated filtered Na^+^ load in diabetes, resulting in Na^+^ excretion comparable to a non-diabetic kidney; see Figs. 2A and 4A1. Due to the coupled action of NKCC2, elevated Na^+^ reabsorption is followed by increases in the reabsorption of Cl^-^(Fig. 2C and 2E). Urine output is predicted to be 30 and 35% higher in moderately diabetic women and men, respectively, due to osmotic diuresis (Figs. 2E and 4A5). Higher filtered K^+^ load enhances its tubular reabsorption, similar to Na^+^, along the proximal tubules and thick ascending limbs (Fig. 2B). These competing factors result in kaliuresis, with K^+^ excretion predicted to be 78 and 48% higher in women and men with moderate diabetes respectively (Figs. 2B and 4A2).

Severe diabetes induces GFR and filtered solute loads by 24% above baseline. The resulting changes in tubular transport are similar to moderate diabetes. In both women and men, the enhanced Na^+^ transport along the proximal tubules and thick ascending limbs essentially compensates for the elevated filtered Na^+^ load in diabetes to limit natriuresis (Figs. 2A and 4A1). Osmotic diuresis approximately doubles urine output (Figs. 2E and 4A5), and urinary K^+^ excretion is approximately two-third higher (Figs. 2B and 4A2).

In sum, diabetes markedly increases GFR and filtered solute loads. That is accompanied by tubular hypertrophy and enhanced renal transport capacity, which prevents excessive loss in fluid and electrolytes (Fig. 4A). The model predicts that the kidneys of women and men with diabetes handle water, electrolytes, and glucose in manners that are qualitatively similar. In the next set of simulations, we examine sex differences in kidney function with SGLT2 inhibition.

### 3.2 Kidney function in non-diabetic women and men under SGLT2 inhibition

The model simulates the administration of a SGLT2 blockade by inhibiting 90% of SGLT2. In a non-diabetic kidney, the remaining 10% of the SGLT2 mediates the reabsorption of 28 and 26% of the filtered glucose along the proximal convoluted tubules in women and men, respectively. The much-elevated glucose flow into the S3 segments increases that segment’s SGLT1-mediated glucose transport to 39 and 43% of filtered load in women and men, respectively. Due to its osmotic diuretic effect of the SGLT2 inhibition [27, 35], passive paracellular reabsorption is attenuated in the proximal tubule. The higher luminal flow stimulates active transport (via torque-induced increases in transcellular transporter expression [46]), but the reduction in passive transport dominates. As a result, net Na^+^ reabsorption decreases in the proximal tubules, by 9.1 and 7.9% in women and men respectively. Solute transport shifts downstream, primarily to the medullary thick ascending limbs, but also to the distal segments (Fig. 5A). The increase in thick ascending limb Na^+^ transport is larger in women compared to men, resulting in more severe natriuresis in men, who exhibit a 265% increase in Na^+^ excretion with SGLT2 inhibition, compared to a 46% increase in women. The elevated Na^+^ reabsorption along the connecting tubules is accompanied by enhanced K^+^ secretion (Fig. 5B). Despite the enhanced transport, SGLT2 inhibition in a non-diabetic kidney induces diuresis and kaliuresis. Urine output increases by 76% in women and to an even larger extent in men by 177%. Similarly, K^+^ excretion increases by 82% and 164% in women and men, respectively (Fig. 4, panels B1–B5).

**Figure 5.**
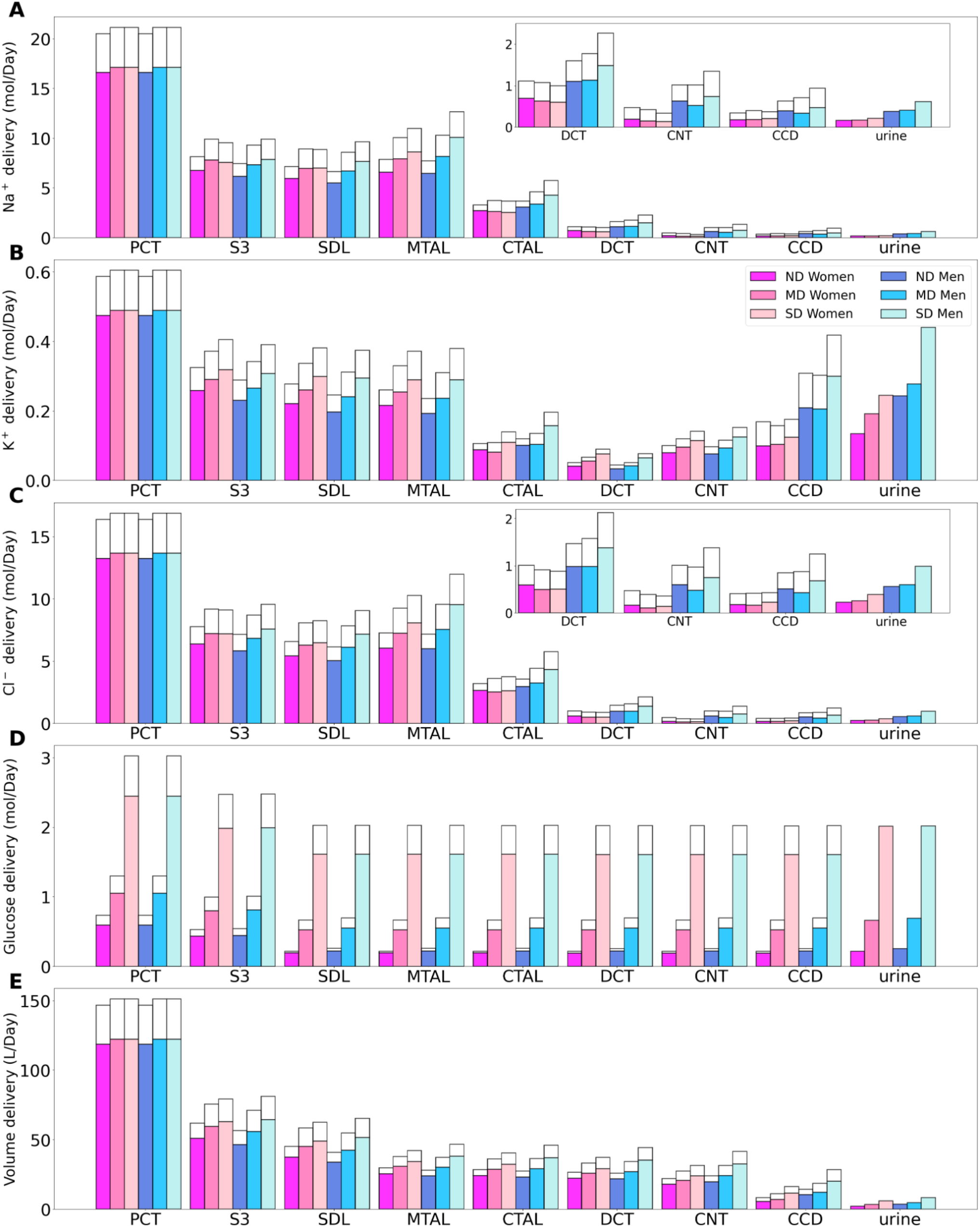
Delivery of Na^+^ (*A*), K^+^ (*B*), Cl^-^ (*C*), glucose (*D*) and fluid (*E*) to the beginning of individual nephron segments in women and men without diabetes (ND), with moderate diabetes (MD), and with severe diabetes (SD) under SGLT2 inhibition. Notations are analogous to Fig. 2.

Taken together, SGLT2 inhibition not only induces glycosuria, but also diuresis, natriuresis, and kaliuresis in non-diabetic men and women. Unlike the earlier results without SGLT2 inhibition, these SGLT2 inhibition results show significant sex differences. The effects on urine excretion are stronger in men, in part due to the larger Na^+^ transport capacity of the thick ascending limbs in the kidneys of women, which allow those segments to compensate for the reduction in transport upstream.

### 3.3 Kidney function in diabetic women and men under SGLT2 inhibition

Kidney function under SGLT2 inhibition is arguably the most clinically relevant case. We would like to recall that SGLT2 inhibition attenuates diabetes-induced glomerular hyperfiltration and returns GFR to baseline, lowering the filtered glucose load from 1.52 to 1.3 mol·day^−1^ in moderate diabetes. The kidney’s response in glucose transport is similar in women and men: proximal convoluted tubule glucose reabsorption, mediated by the 10% remaining SGLT2, reduces by ~80%, from 1.5 to 0.3 mol glucose·day^−1^. The SGLT1 along the S3 segment reabsorbs a fraction of the remaining glucose, at a rate similar to the proximal convoluted tubule, thereby limiting the risk of hypoglycemia [47]. Glucose excretion in moderately diabetic women and men is predicted to be similar, at 0.69 and 0.66 mol·day^−1^, respectively, which corresponds to almost 50% of filtered glucose; see Fig. 4B4.

The kidney’s response to SGLT2 inhibition in severe diabetes exhibits similarities but also notable differences from the moderate diabetes case. Even though SGLT2 inhibition normalizes GFR, the plasma glucose level remains high at 20 mM, which yields a filtered glucose load of 3.0 mol glucose·day^−1^. Proximal tubule glucose transport is similar in women and men with severe diabetes, with the proximal convoluted tubule and S3 reabsorbing ~0.55 and 0.45 mol glucose·day^−1^, respectively (Fig. 6D). Together, these two segments account for two-third of the filtered glucose. The predicted glucose excretion of 2.02 mol·day^−1^ in both sexes (Fig. 4B4) is consistent with reported values [48]. A major difference between the moderate and severe diabetes cases lies in the S3 response. Recall that in moderate diabetes, glucose transport along the S3 segment jumps from negligible to ~0.3 mol glucose.day^-1^, which corresponds to 23% of filtered glucose following SGLT2 inhibition. In contrast, in severe diabetes, even without SGLT2 inhibition, S3 already mediates the reabsorption of a significant fraction of the filtered glucose (0.36 mol glucose.day^-1^ or 9% of the filtered glucose), because the glucose transport capacity of the proximal convoluted tubule is overwhelmed. Following SGLT2 inhibition, the increase in S3 glucose transport is less drastic, specifically, an increase of 25% to 0.45 mol glucose.day^-1^ (compare Figs. 4A4 and 4B4). Also, whereas there is glucose secretion across the tight junction of the proximal convoluted tubules in the absence of treatment, the paracellular pathway mediates glucose reabsorption when SGLT2 is inhibited (owing to higher luminal glucose concentration).

**Figure 6.**
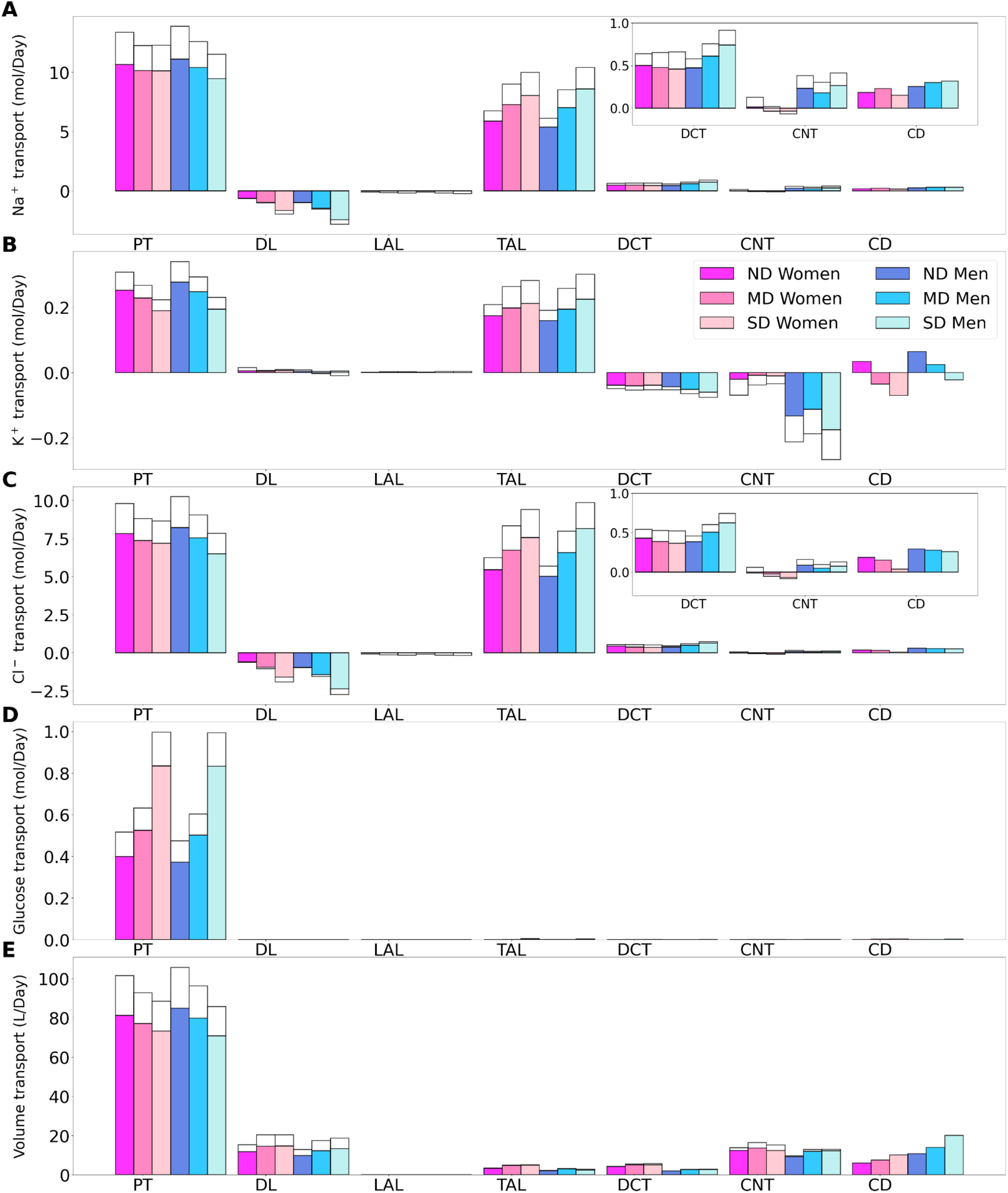
Transport of Na^+^ (*A*), K^+^ (*B*), Cl^-^ (*C*), glucose (*D*) and fluid (*E*) along individual nephron segments in women and men without diabetes (ND), with moderate diabetes (MD), and with severe diabetes (SD) under SGLT2 inhibition. Notations are analogous to figure 2. PT, proximal convoluted tubule and S3.

While the primary target of SGLT2 inhibitors is glucose, they also have a significant impact on renal Na^+^ transport. By normalizing GFR, SGLT2 inhibition significantly lowers filtered Na^+^ load and Na^+^ transport. Our simulations suggest that, SGLT2 inhibition induces osmotic diuresis in the proximal tubules, as in the non-diabetic case, thereby reducing paracellular transport. In fact, the model predicts that osmotic diuresis reverses the direction of paracellular Na^+^ transport in S3: the luminal-to-interstitial Na^+^ concentration gradient favors Na^+^ secretion into the lumen via the tight junctions. Consequently, in the moderate diabetes case, Na^+^ excretion increases by 68 and 228% in women and men, respectively (Fig. 4B1). Natriuresis is less severe in women because the thick ascending limbs in their kidneys are more capable of compensating for the reduced Na^+^ transport by the proximal convoluted tubules. Importantly, natriuresis is also accompanied by diuresis, with urine output increases of 109 and 170%, in women and men, respectively. The elevated Na^+^ along the distal tubular segments raises K^+^ secretion. In men, that yields an increase of 103% in K^+^ excretion compared to the cases without SGLT2 inhibition (Fig. 4B2). In women, the distal tubular Na^+^ flow is lower than in men, as is K^+^ secretion. As such, kaliuresis is attenuated in women, with K^+^ excretion 45% higher than without SGLT2 inhibition. Qualitatively similar results are obtained for the severe diabetes case; see Figs. 4B1, 5A and 6A.

Taken together our results indicate that the intended glycosuric effect of SGLT2 inhibitors is similar in women and men with diabetes. SGLT2 inhibition also induces diuresis, natriuresis, and kaliuresis, to a more severe extent in men compared to women.

### 3.4 The influence of sex and SGLT2 inhibition on the tubuloglomerular feedback signal

At the onset of diabetes, the kidney hypertrophies and SNGFR increases. A causal link between the two pathophysiological processes has been proposed, based on the tubuloglomerular feedback. The tubuloglomerular feedback adjusts SNGFR based on the luminal [Cl^−^] sensed by the macula densa cells: if the [Cl^−^] exceeds a target value, the tubuloglomerular feedback is activated, and a signal is sent to constrict the afferent arteriole and reduce SNGFR, and vice versa [49]. The tubuloglomerular feedback is believed to play a role in the diabetes-induced glomerular hyperfiltration: the enhanced proximal tubular transport lowers the tubuloglomerular feedback signal, i.e., the luminal [Cl^−^] sensed by the macula densa cells, resulting in a feedback-induced increase in SNGFR [50]. Because the tubuloglomerular feedback is not characterized in humans, it is not explicitly represented in the present models. Instead, SNGFR is assumed to be known a priori. Nevertheless, we present here our model results to assess the potential influences of sex and SGLT2 inhibition on the tubuloglomerular feedback, and the implications on SNGFR.

Table 2 shows predicted luminal [Cl^-^] at the macula densa, where the sensing of the tubuloglomerular feedback signal occurs and is taken to be the end of the cortical thick ascending limbs. Values are computed for the superficial nephron, and as weighted averages of the juxtamedullary nephrons. For both women and men, diabetes lowers macula densa [Cl^-^], which would attenuate the tubuloglomerular feedback signal and would lead to glomerular hyperfiltration. The macula densa [Cl^-^] is predicted to be lower in women in all cases. Taken in isolation, a lower macula densa [Cl^-^] should correspond to higher SNGFR. In contrast, after correction for weight, GFR is not known to differ significantly between sexes [51, 52]. Thus, the kidneys of women and men may have different tubuloglomerular feedback operating points. That is, given different baseline macula densa [Cl^-^] values, the different tubuloglomerular feedback systems in women and men would generate similar SNGFR.

**Table 2:**
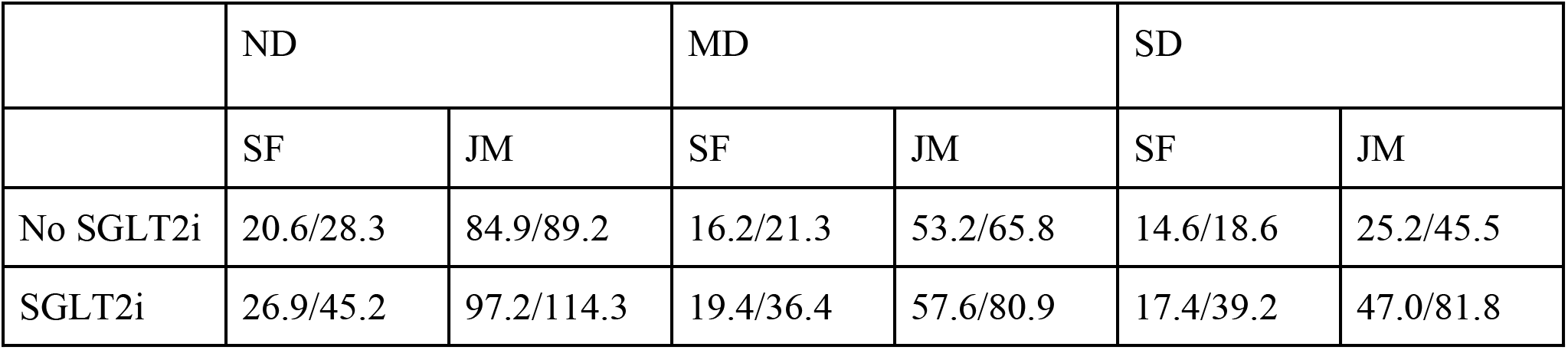
Luminal [Cl^-^] at the macula densa, given as [female value]/[male value]. SGLT2i, SGLT2 inhibition; ND, no diabetes; MD, moderate diabetes; SD, severe diabetes; SF, superficial nephron; JM, weighted average of model juxtamedullary nephrons.

|  | ND |  | MD |  | SD |  |
| --- | --- | --- | --- | --- | --- | --- |
|  | SF | JM | SF | JM | SF | JM |
| No SGLT2i | 20.6/28.3 | 84.9/89.2 | 16.2/21.3 | 53.2/65.8 | 14.6/18.6 | 25.2/45.5 |
| SGLT2i | 26.9/45.2 | 97.2/114.3 | 19.4/36.4 | 57.6/80.9 | 17.4/39.2 | 47.0/81.8 |

The model predictions also indicate a GFR-normalizing effect of SGLT2 inhibition. By limiting Na^+^-glucose transport, SGLT2 inhibition increases macula densa [Cl^-^] substantially (Table 2), which would activate the tubuloglomerular feedback and lower SNGFR. It is noteworthy that the increases in macula densa [Cl^-^] are significantly larger in men than in women. Taken in isolation, that might suggest that SGLT2 inhibition may be more effective in ameliorating diabetes-induced glomerular hyperfiltration in men compared to women. However, such sex difference has not been reported, a discrepancy that may be attributed to the involvement of factors not represented in the model (see Discussion).

## 4 Discussion

In addition to regulating fluid, electrolytes, and blood pressure, the kidney also plays a key role in glucose homeostasis. In the euglycemic state, essentially all the filtered glucose is reabsorbed by the proximal tubules via the SGLT2 and SGLT1, with little glucose excreted in urine. In diabetes, where either insulin production or sensitivity is impaired, the blood glucose level rises. Chronic exposure to elevated blood glucose levels contribute to the tubular hypertrophy observed in diabetes. In turn, this increase in epithelial cell area due to tubular hypertrophy increases the transport capacity, which may increase SNGFR by suppressing tubuloglomerular feedback (more below). The activities of SGLT2/SGLT1 are likely modified in patients with diabetes; however, those changes have not been characterized. Nonetheless, when sufficiently high plasma glucose is coupled with glomerular hyperfiltration, the kidney’s glucose transport capacity may be overwhelmed, resulting in the appearance of glucose in urine, which is traditionally considered a hallmark of diabetes. In fact, the Chinese translation of “diabetes” is literally “sugar in urine” disease.

The model findings presented here have provided insights into the mechanisms that given rise to glomerular hyperfiltration in early diabetes, its normalization by SGLT2 inhibitors, and potential sex differences. Consistent with the “tubulo-centric” hypothesis by Vallon and co-workers, predictions by both the models for women and men indicate that the enhanced reabsorption along the proximal tubules in early diabetes lowers the luminal [Cl^−^] sensed by the macula densa cells, suppresses the tubuloglomerular feedback signal, and results in a feedback-induced increase in SNGFR. These model findings are consistent with micropuncture findings in rats. It must be noted that the presence of a tubuloglomerular feedback has yet to be established in the human kidney. Nonetheless, if tubuloglomerular feedback is present in humans, the vasodilative signal predicted in diabetes could contribute to glomerular hyperfiltration. In addition, if humans exhibit the sexual dimorphism in renal transporter pattern as observed in rodents, then the tubuloglomerular feedback signal likely differs between women and men. For women and men to have similar SNGFR, the operating points of their tubuloglomerular feedback systems may have to be adjusted accordingly, to map different macula densa [Cl^-^] values to the same SNGFR.

Moreover, the model predicts that SGLT2 inhibition while lowering Na^+^ and glucose transport, significantly increases luminal [Cl^−^] at the macula densa (Table 1), which may then attenuate glomerular hyperfiltration via tubuloglomerular feedback, consistent with clinical data, which show a reduction in eGFR in patients with diabetes following chronic SGLT2 inhibition. The model predicts that SGLT2 inhibition increases macula densa luminal [Cl^−^] to a larger extent in men than in women. However, such sex difference in drug response has not been reported. That discrepancy may be attributed to factors such as the resetting of tubuloglomerular feedback (i.e., adjustment of its operating point) [53], or sex-specific transporter alterations in diabetes and SGLT2 inhibition that have yet to be characterized. Nevertheless, for both women and men, lowering glomerular hyperfiltration on the single nephron level by SGLT2 inhibition may provide long-term beneficial effects in the diabetic kidney, by reducing the transport load and metabolic requirement on the nephrons [31, 54].

In sum, we have developed the sex-specific computational models of detailed epithelial transport in the kidney of a patient with diabetes. The models predict that, similar to rodents, diabetes-induced tubular hypertrophy in both men and women may contribute to glomerular hyperfiltration via a (hypothesized) tubuloglomerular feedback signal. Model predictions further suggest that SGLT2 inhibition may activate tubuloglomerular feedback and attenuate glomerular hyperfiltration, and the tubuloglomerular feedback signal may be stronger in men. Model simulations did not reveal significant differences in renal handling of electrolyte and water handling between women and men with diabetes. More notable sex differences emerge in simulations of SGLT2 inhibition, which induces diuresis, natriuresis, and kaliuresis, with those effects significantly more prominent in men compared to women. In both men and women, the predicted increases in urine output and excretion are significantly higher than experimentally reported values [55]. For instance, the model predicts that in the moderate diabetes case, urine output doubles in women, and almost triples in men (Fig. 4B). In contrast, empagliflozin increases urine volume by 24% [55]. Excessive urine output may result in volume depletion, which in turn activates mechanisms to reduce urine production. Because most of these mechanisms are external to the kidney, they are not represented in the present model, which may explain the excessive urine output and excretion predicted by the model.

A major limitation of the present models is that the interstitial fluid composition is assumed to be known a priori. As such, the models do not capture the interactions among different nephron segments. A worthwhile extension is to incorporate these interactions and perhaps capture the spatially heterogeneous distribution of the renal tubules [56–58]. Moreover, as previously noted, tubuloglomerular feedback is not explicitly represented. Once data describing that feedback system in humans become available, the present model can be combined with a model of the tubuloglomerular feedback [59, 60], so that changes in SNGFR following drug treatments can be predicted. Besides controlling solute transport, the kidney also filters and secretes uremic toxins such as indoxyl sulfate, hippuric acid and p-cresyl sulfate, important food and drug organic metabolites whose accumulation the blood induces many complications and ultimately lead to kidney failure [61–63]. Importantly, the uremic toxin transporters such as organic anionic transporter (OAT) 1 and 3 also display sex-differences [12–14]. Moreover, evidence shows that the OAT3 expression is reduced in diabetic rats [64, 65], and as such including uremic toxins and their transporters represents an important avenue for model extension. Despite its limitations, the present models are a stepping stone to evaluate the renal mechanisms under the effect of commonly-prescribed medications besides SGLT2 inhibitors, such as blockers of the angiotensin II system, in both women and men with diabetes, and to assess the the impact of impaired kidney function.

## 6 Author Contributions

Conceptualization, A.T.L..; methodology, A.T.L.; software, A.T.L.; validation, A.T.L. and S.S.; formal analysis, A.T.L. and S.S.; investigation, A.T.L. and S.S.; resources, A.T.L.; data curation, A.T.L.; writing—original draft preparation, A.T.L., S.S. and A.C.; writing—review and editing, A.T.L., S.S. and A.C.; visualization, S.S.; supervision, A.T.L.; project administration, A.T.L.; funding acquisition, A.T.L. All authors have read and agreed to the published version of the manuscript.

## 8 Acknowledgments

None

## 9 Supplementary Material

Supplementary Material should be uploaded separately on submission, if there are Supplementary Figures, please include the caption in the same file as the figure. Supplementary Material templates can be found in the Frontiers Word Templates file.

Please see the <u>Supplementary Material section of the Author guidelines</u> for details on the different file types accepted.

## 10 Data Availability Statement

Computer code developed for this study is available at https://github.com/uwrhu/Python-nephron-model-parallel-code-latest

## References

1. Saeedi, P., et al., Global and regional diabetes prevalence estimates for 2019 and projections for 2030 and 2045: Results from the International Diabetes Federation Diabetes Atlas. Diabetes research and clinical practice, 2019. 157: p. 107843.

2. Foley, R.N. and A.J. Collins, End-stage renal disease in the United States: an update from the United States Renal Data System. Journal of the American Society of Nephrology, 2007. 18(10): p. 2644–2648.

3. Koye, D.N., et al., The global epidemiology of diabetes and kidney disease. Advances in chronic kidney disease, 2018. 25(2): p. 121–132.

4. Bjornstad, P. and D.Z. Cherney, Renal hyperfiltration in adolescents with type 2 diabetes: physiology, sex differences, and implications for diabetic kidney disease. Current diabetes reports, 2018. 18(5): p. 1–7.

5. Lovshin, J.A., et al., Hyperfiltration, urinary albumin excretion, and ambulatory blood pressure in adolescents with type 1 diabetes mellitus. American Journal of Physiology-Renal Physiology, 2018. 314(4): p. F667–F674.

6. Hatano, R., et al., Sex hormones induce a gender-related difference in renal expression of a novel prostaglandin transporter, OAT-PG, influencing basal PGE2 concentration. American Journal of Physiology-Renal Physiology, 2012. 302(3): p. F342–F349.

7. Hilliard, L.M., et al., Gender differences in pressure-natriuresis and renal autoregulation: role of the angiotensin type 2 receptor. Hypertension, 2011. 57(2): p. 275–282.

8. Reckelhoff, J.F., Gender differences in the regulation of blood pressure. Hypertension, 2001. 37(5): p. 1199–1208.

9. Sabolić, I., et al., Gender differences in kidney function. Pflügers Archiv-European Journal of Physiology, 2007. 455(3): p. 397–429.

10. Sullivan, J.C. and E.E. Gillis, Sex and gender differences in hypertensive kidney injury. American Journal of Physiology-Renal Physiology, 2017. 313(4): p. F1009–F1017.

11. Veiras, L.C., et al., Sexual dimorphic pattern of renal transporters and electrolyte homeostasis. Journal of the American Society of Nephrology, 2017. 28(12): p. 3504–3517.

12. Urakami, Y., et al., Gender differences in expression of organic cation transporter OCT2 in rat kidney. FEBS letters, 1999. 461(3): p. 339–342.

13. Groves, C.E., et al., Sex differences in the mRNA, protein, and functional expression of organic anion transporter (Oat) 1, Oat3, and organic cation transporter (Oct) 2 in rabbit renal proximal tubules. Journal of Pharmacology and Experimental Therapeutics, 2006. 316(2): p. 743–752.

14. Euteneuer, A.M., et al., Estrogen receptor a (ERa) indirectly induces transcription of human renal organic anion transporter 1 (OAT1). Physiological reports, 2019. 7(21): p. e14229.

15. Breljak, D., et al., Sex-dependent expression of Oat3 (Slc22a8) and Oat1 (Slc22a6) proteins in murine kidneys. American Journal of Physiology-Renal Physiology, 2013. 304(8): p. F1114–F1126.

16. Hu, R., A.A. McDonough, and A.T. Layton, Functional implications of the sex differences in transporter abundance along the rat nephron: modeling and analysis. American Journal of Physiology-Renal Physiology, 2019. 317(6): p. F1462–F1474.

17. Hu, R., A.A. McDonough, and A.T. Layton, Sex differences in solute transport along the nephrons: effects of Na+ transport inhibition. American Journal of Physiology-Renal Physiology, 2020. 319(3): p. F487–F505.

18. Li, Q., et al., Functional implications of sexual dimorphism of transporter patterns along the rat proximal tubule: modeling and analysis. American Journal of Physiology-Renal Physiology, 2018. 315(3): p. F692–F700.

19. Chen, Y., et al., Sex-specific computational models of the spontaneously hypertensive rat kidneys: factors affecting nitric oxide bioavailability. American Journal of Physiology-Renal Physiology, 2017. 313(2): p. F174–F183.

20. Fry, B.C., A. Edwards, and A.T. Layton, Impacts of nitric oxide and superoxide on renal medullary oxygen transport and urine concentration. American Journal of Physiology-Renal Physiology, 2015. 308(9): p. F967–F980.

21. Fry, B.C., A. Edwards, and A.T. Layton, Impact of nitric-oxide-mediated vasodilation and oxidative stress on renal medullary oxygenation: a modeling study. American Journal of Physiology-Renal Physiology, 2016. 310(3): p. F237–F247.

22. Fry, B.C., et al., Impact of renal medullary three-dimensional architecture on oxygen transport. American Journal of Physiology-Renal Physiology, 2014. 307(3): p. F263–F272.

23. Chen, J., A. Edwards, and A.T. Layton, Effects of pH and medullary blood flow on oxygen transport and sodium reabsorption in the rat outer medulla. American Journal of Physiology-Renal Physiology, 2010. 298(6): p. F1369–F1383.

24. Hu, R., A.A. McDonough, and A.T. Layton, Sex differences in solute and water handling in the human kidney: Modeling and functional implications. iScience, 2021: p. 102667.

25. Maric-Bilkan, C. Sex differences in diabetic kidney disease. Elsevier.

26. Mogensen, C.E., Glomerular filtration rate and renal plasma flow in short-term and longterm juvenile diabetes mellitus. Scandinavian journal of clinical and laboratory investigation, 1971. 28(1): p. 91–100.

27. Layton, A.T., V. Vallon, and A. Edwards, Predicted consequences of diabetes and SGLT inhibition on transport and oxygen consumption along a rat nephron. American Journal of Physiology-Renal Physiology, 2016. 310(11): p. F1269–F1283.

28. Shepard, B.D., Sex differences in diabetes and kidney disease: mechanisms and consequences. American Journal of Physiology-Renal Physiology, 2019. 317(2): p. F456–F462.

29. Chao, E.C. and R.R. Henry, SGLT2 inhibition—a novel strategy for diabetes treatment. Nature reviews drug discovery, 2010. 9(7): p. 551–559.

30. Carpentier, C., et al., Glycosuria amount in response to hyperglycaemia and risk for diabetic kidney disease and related events in Type 1 diabetic patients. Nephrology dialysis transplantation, 2019. 34(10): p. 1731–1738.

31. Neal, B., et al., Canagliflozin and cardiovascular and renal events in type 2 diabetes. New England Journal of Medicine, 2017. 377(7): p. 644–657.

32. Zinman, B., et al., Empagliflozin, cardiovascular outcomes, and mortality in type 2 diabetes. New England Journal of Medicine, 2015. 373(22): p. 2117–2128.

33. Hu, R. and A. Layton, A computational model of kidney function in a patient with diabetes. International Journal of Molecular Sciences, 2021. 22(11): p. 5819.

34. Layton, A.T. and V. Vallon, SGLT2 inhibition in a kidney with reduced nephron number: modeling and analysis of solute transport and metabolism. American Journal of Physiology-Renal Physiology, 2018. 314(5): p. F969–F984.

35. Layton, A.T., V. Vallon, and A. Edwards, Modeling oxygen consumption in the proximal tubule: effects of NHE and SGLT2 inhibition. American Journal of Physiology-Renal Physiology, 2015. 308(12): p. F1343–F1357.

36. Layton, A.T. and H.E. Layton, A computational model of epithelial solute and water transport along a human nephron. PLoS computational biology, 2019. 15(2): p. e1006108.

37. Oliver, J. Nephrons and kidneys: a quantitative study of developmental and evolutionary mammalian renal architectonics. 1968.

38. Layton, A.T., et al., Solute transport and oxygen consumption along the nephrons: effects of Na+ transport inhibitors. American Journal of Physiology-Renal Physiology, 2016. 311(6): p. F1217–F1229.

39. Layton, A.T., V. Vallon, and A. Edwards, A computational model for simulating solute transport and oxygen consumption along the nephrons. American Journal of Physiology-Renal Physiology, 2016. 311(6): p. F1378–F1390.

40. Sabolić, I., et al., Expression of Na+-D-glucose cotransporter SGLT2 in rodents is kidney-specific and exhibits sex and species differences. American Journal of Physiology-Cell Physiology, 2012. 302(8): p. C1174–C1188.

41. Nauck, M.A., Update on developments with SGLT2 inhibitors in the management of type 2 diabetes. Drug design, development and therapy, 2014. 8: p. 1335.

42. Baumgartl, H.-J., et al., On the prognosis of IDDM patients with large kidneys. Nephrology, dialysis, transplantation: official publication of the European Dialysis and Transplant Association-European Renal Association, 1998. 13(3): p. 630–634.

43. Seman, L., et al., Empagliflozin (BI 10773), a potent and selective SGLT2 inhibitor, induces dose-dependent glucosuria in healthy subjects. Clinical pharmacology in drug development, 2013. 2(2): p. 152–161.

44. Cherney, D.Z.I., et al., Renal hemodynamic effect of sodium-glucose cotransporter 2 inhibition in patients with type 1 diabetes mellitus. Circulation, 2014. 129(5): p. 587–597.

45. Edwards, A., et al., Effects of NKCC2 isoform regulation on NaCl transport in thick ascending limb and macula densa: a modeling study. Am J Physiol Renal Physiol, 2014. 307(2): p. F137–46.

46. Layton, A.T., A. Edwards, and V. Vallon, Adaptive changes in GFR, tubular morphology, and transport in subtotal nephrectomized kidneys: modeling and analysis. American Journal of Physiology-Renal Physiology, 2017. 313(2): p. F199–F209.

47. Nespoux, J. and V. Vallon, SGLT2 inhibition and kidney protection. Clinical Science, 2018. 132(12): p. 1329–1339.

48. List, J.F. and J.M. Whaley, Glucose dynamics and mechanistic implications of SGLT2 inhibitors in animals and humans. Kidney International, 2011. 79: p. S20–S27.

49. Layton, A.T., Feedback-mediated dynamics in a model of a compliant thick ascending limb. Mathematical biosciences, 2010. 228(2): p. 185–194.

50. Sgouralis, I. and A.T. Layton, Theoretical assessment of renal autoregulatory mechanisms. American Journal of Physiology-Renal Physiology, 2014. 306(11): p. F1357–F1371.

51. de Hauteclocque, A., et al., Complications The influence of sex on renal function decline in people with Type 2 diabetes. 2014.

52. Jacobsen, P., et al., Progression of diabetic nephropathy in normotensive type 1 diabetic patients. Kidney International, 1999. 56: p. S101–S105.

53. Thomson, S.C. and R.C. Blantz, Glomerulotubular balance, tubuloglomerular feedback, and salt homeostasis. Journal of the American Society of Nephrology, 2008. 19(12): p. 2272–2275.

54. Ferrannini, E. and A. Solini, SGLT2 inhibition in diabetes mellitus: rationale and clinical prospects. Nature Reviews Endocrinology, 2012. 8(8): p. 495–502.

55. Mordi, N.A., et al., Renal and cardiovascular effects of SGLT2 inhibition in combination with loop diuretics in patients with type 2 diabetes and chronic heart failure: the RECEDE-CHF trial. Circulation, 2020. 142(18): p. 1713–1724.

56. Pannabecker, T.L., et al., Role of three-dimensional architecture in the urine concentrating mechanism of the rat renal inner medulla. American Journal of Physiology-Renal Physiology, 2008. 295(5): p. F1271–F1285.

57. Layton, A.T., W.H. Dantzler, and T.L. Pannabecker, Urine concentrating mechanism: impact of vascular and tubular architecture and a proposed descending limb urea-Na+ cotransporter. American Journal of Physiology-Renal Physiology, 2012. 302(5): p. F591–F605.

58. Pannabecker, T.L. and A.T. Layton, Targeted delivery of solutes and oxygen in the renal medulla: role of microvessel architecture. American Journal of Physiology-Renal Physiology, 2014. 307(6): p. F649–F655.

59. Chen, J., et al., A mathematical model of the myogenic response to systolic pressure in the afferent arteriole. American Journal of Physiology-Renal Physiology, 2011. 300(3): p. F669–F681.

60. Sgouralis, I., et al., Renal hemodynamics, function, and oxygenation during cardiac surgery performed on cardiopulmonary bypass: a modeling study. Physiological reports, 2015. 3(1): p. e12260.

61. Duranton, F., et al., Normal and pathologic concentrations of uremic toxins. Journal of the American Society of Nephrology, 2012. 23(7): p. 1258–1270.

62. Herget-Rosenthal, S., et al. Progress in uremic toxin research: uremic toxins in acute kidney injury. Wiley Online Library.

63. Schophuizen, C.M.S., et al., Cationic uremic toxins affect human renal proximal tubule cell functioning through interaction with the organic cation transporter. Pflügers Archiv-European Journal of Physiology, 2013. 465(12): p. 1701–1714.

64. Lungkaphin, A., et al., Impaired insulin signaling affects renal organic anion transporter 3 (Oat3) function in streptozotocin-induced diabetic rats. PLoS One, 2014. 9(5): p. e96236.

65. Phatchawan, A., et al., Decreased renal organic anion transporter 3 expression in type 1 diabetic rats. The American Journal of the Medical Sciences, 2014. 347(3): p. 221–227.

